# Voluntary control of human motor units is more flexible in facial than limb muscles

**DOI:** 10.64898/2026.11.21.767708

**Authors:** Eric A. Kirk, Sophia T. Jenz, James A. Beauchamp, Alex T. Benedetto, Charles J. Heckman, Gregory E.P. Pearcey, Charles L. Rice

**Author notes:** E. Kirk and S. Jenz made equal contributions as first authors. G. Pearcey and C. Rice made equal contributions as senior authors. Correspondence Eric Kirk.

## Abstract

Facial expressions require different motor circuits than those controlling the limbs. How the motor system uses flexible neural control across muscles innervated by the brainstem versus the spinal cord is unclear. Using intramuscular and surface electromyography in human participants performing low-effort, sustained voluntary contractions, we show that brainstem motor units function with more features of flexible control compared to motor units in several spinally innervated muscles. Individual motor units recorded from the frontalis muscle (brainstem-innervated) used a wider range of regular and irregular firing rates, reflecting both greater independence and higher input sensitivity than spinal-innervated motor units. When assessing motor unit pairs, we again found that brainstem control relied more heavily on features corresponding to independent inputs relative to spinal control. Voluntary control of brainstem-innervated muscle therefore used more flexible neural strategies than the control of upper and lower limb muscles we tested. These findings support an anatomical hypothesis for adaptive neural control, in which the degree of independent motor unit activation is constrained by how muscles are accessed across cortex.

## Introduction

The motor system integrates a large amount of information to control muscles that maintain posture and express emotions. Muscles used in facial expressions have evolved separate innervation circuits and rely more heavily on cortical inputs^1–7^ than those that serve the limbs^8,9^. This suggests that these disparate motor pools require different rules for voluntary control. For example, muscles of the upper face are often used as a diagnostic criterion when assessing peripheral versus central palsy, because people with focal cortical stroke can still wrinkle their forehead as these muscles are governed by both hemispheres^6^. Recent studies characterizing brainstem and spinal motor neurons propose that smaller cell size and different classes of ion channels explain what makes brainstem neurons more excitable^10,11^. However, how voluntary commands are orchestrated between brainstem and spinal motor pools of the central nervous system is poorly understood.

Despite anatomical differences, the neural control of movement ultimately occurs through the activation of motor units. Each motor unit corresponds to one motor neuron residing in the brainstem or spinal cord that innervates a few to many hundred skeletal muscle fibres in the periphery. During voluntary contractions, the majority of evidence suggests that there is rigid control of motor neurons governed by common synaptic inputs and size-related orderly recruitment^12–20^. A recent study, however, in a non-human primate has provided convincing evidence that questions whether the ordered recruitment of motor units is truly rigid by showing properties of flexible synaptic drive in spinally-innervated muscles^21^. For a muscle to operate in a flexible regime, this would mean that motor units would be under independent circuit control that can be accessed from different elements of the cortex, descending tracts, or reflex pathways. Under this assumption, an anatomically based theory of flexible control would be relevant, because some brainstem innervated muscles have strong connections to both cortical hemispheres, as compared to spinally innervated muscles.

A complementary, but often overlooked component of voluntary control involves the firing regime in which the motor unit operates^22^. The contrast between firing output that is more variable, as compared to regular, provides mechanistic understanding for how synaptic inputs are distributed across the motor pool, that drive motor neuron outputs relative to their critical membrane threshold^23–30^. Furthermore, movement features will change depending on which firing regime is used, because muscles have catch-like properties in response to neuronal inputs. For example, low-threshold motor units that have longer contractile relaxation would better integrate irregular firing rates^25,26,31–34^. In this context, two motor units, with one operating in an irregular firing regime (also called fluctuation-driven or sub-primary range) and the other operating with a regular firing regime (also called mean-driven or primary range), would presumably have more flexible control properties as compared to the same pair of motor units that are both only active with regular firing. Overall, combining an understanding of the firing regime and interactions across motor units provides the ability to assess properties of synaptic drive with measures of independent neural control.

In human participants, we recorded motor units in brainstem and spinally innervated muscles during low-effort voluntary contractions. First, we made intramuscular recordings to determine the firing regimes used for each muscle. Then, we made high-density surface array recordings to assess the firing properties across simultaneously recorded motor units. Our results provide support for an anatomical theory of flexible control, in which brainstem motor units broadly use both firing regimes and show more properties of independent control as compared with spinally innervated muscles of the upper and lower limb.

## Results

### Task and motor unit recordings

We studied brainstem- and spinal-innervated muscles during sustained voluntary contractions. The task required participants to activate each muscle with relatively low effort, and to maintain that level of activation throughout the contraction (Fig 1a). Visual feedback of the real-time rectified EMG signal was provided during contractions that were maintained for several minutes. It is known that muscle recordings have inherent noise; therefore, we utilized slower-moving increases and decreases of the EMG amplitude relative to the moving baseline, which were representative of changes in mean synaptic drive to the motor pool (Fig 1b). From the continuous voluntary contraction, we segmented whole-muscle activation curves as a way to understand how the motor unit firing response varied across muscles. From intramuscular recordings in the frontalis (brainstem-innervated), and biceps brachii, anconeus and first dorsal interosseous muscles (spinally-innervated), we were able to measure the output of motor neurons with high accuracy, which is an important technical consideration when using metrics that are sensitive to missing action potentials (Fig 1c).

**Figure 1:**
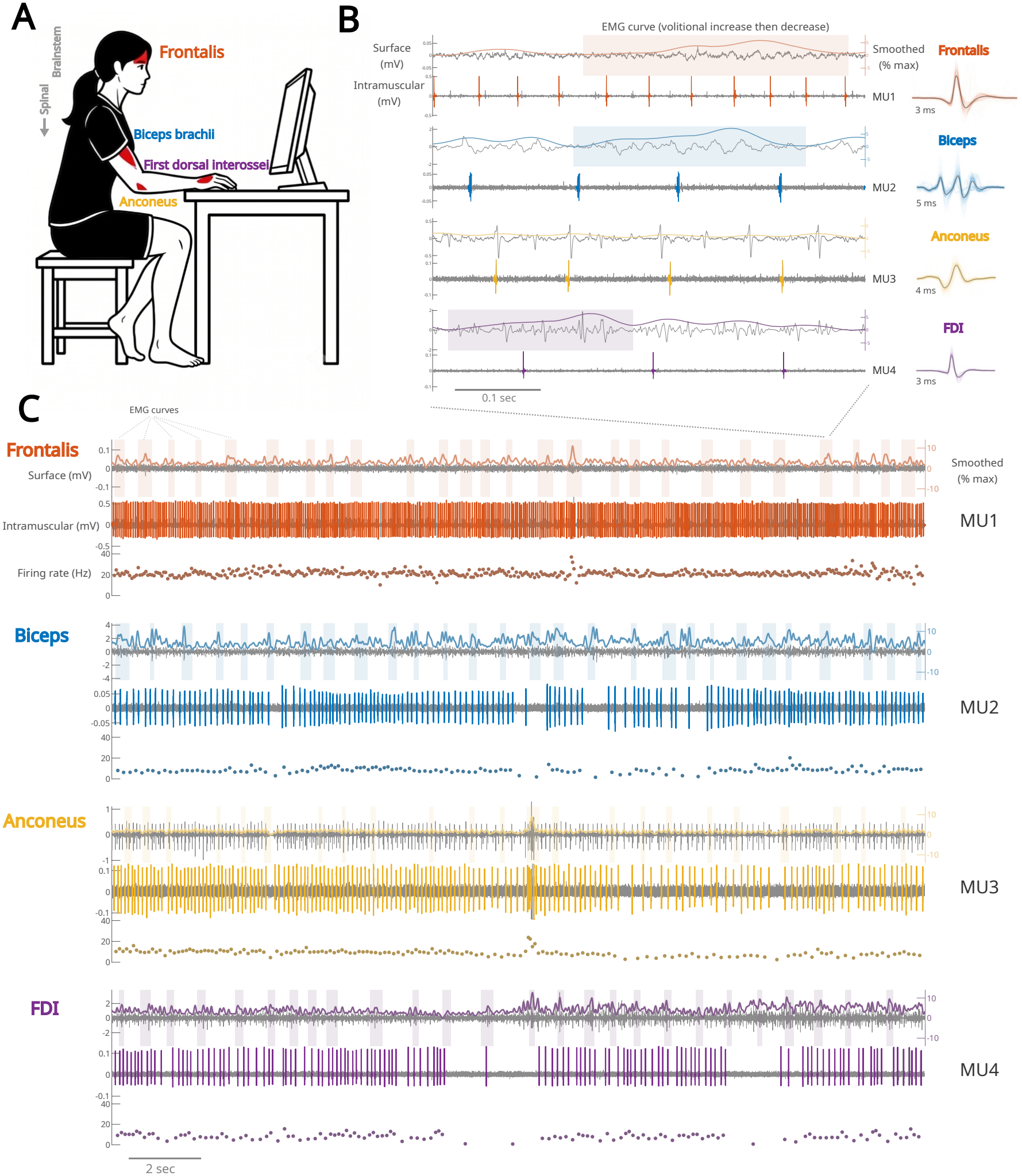
Voluntary activation of facial and upper limb muscles. **A**, cartoon showing the experimental arrangement with the participant seated at a table in a relaxed position, with real-time visual feedback of whole muscle surface electromyography (EMG). We recorded from the right aspect of the frontalis muscle (above the right eye) and from muscles of the right upper limb. Each muscle was recorded independently as the participant attempted to maintain a near-constant voluntary contraction for 3-12 min that was held between 1 and 18% of maximum muscle activation. **B**, 0.5 second recording from each muscle in the next panel. The mean waveform and 150 randomly chosen overlays are shown with trace duration for each motor unit (MU). Highlighted are example surface EMG curves where the averaged muscle activity increased then decreased, used as a proxy for changes of volitional drive to the motor pool. **C**, 20 second recording from each muscle showing the DC-offset-removed surface and intramuscular signals (n = 1 participant). The surface EMG curve colored for each muscle was bandpass filtered, rectified and smoothed (sigma = 100 ms). The colored blocks on the surface recordings show the onset and offset of identified peaks that were used as an indirect estimate of increases and decreases of average synaptic input to the motor pool for each muscle. A single motor unit for each muscle is identified and marked for each intramuscular channel.

During contractions, real-time feedback of surface EMG was used instead of joint-produced torque. This enabled similar comparisons across muscles because it was not possible to record force from the frontalis. Likewise, because contractions required lower levels of effort, this enabled better signal-to-noise due to fewer recruited motor units, and minimized the confound of muscle fatigue. As a consequence, our sample across muscles consisted of lower threshold motor units. Generally, we found that frontalis and first dorsal interosseous motor unit action potential waveforms had a narrow width, whereas waveforms for other spinally innervated muscles were larger and had more complexity (example, Fig 1b).

### Identifying firing regimes between brainstem and spinal motor units

Assessing characteristics of firing rates provides mechanistic understanding about the role of the synaptic inputs, noise, and properties of the transformation into action potentials^26,32,35–38^. Recent work modelling skewness of cortical^24^ and irregularity of spinal firing^23^ has shown how neuronal activity operates across two regimes. These regimes, defined as fluctuation- and mean-driven, are largely dependent on the relationship between the level of synaptic input relative to the membrane voltage threshold for action potentials (Fig 2a).

**Figure 2:**
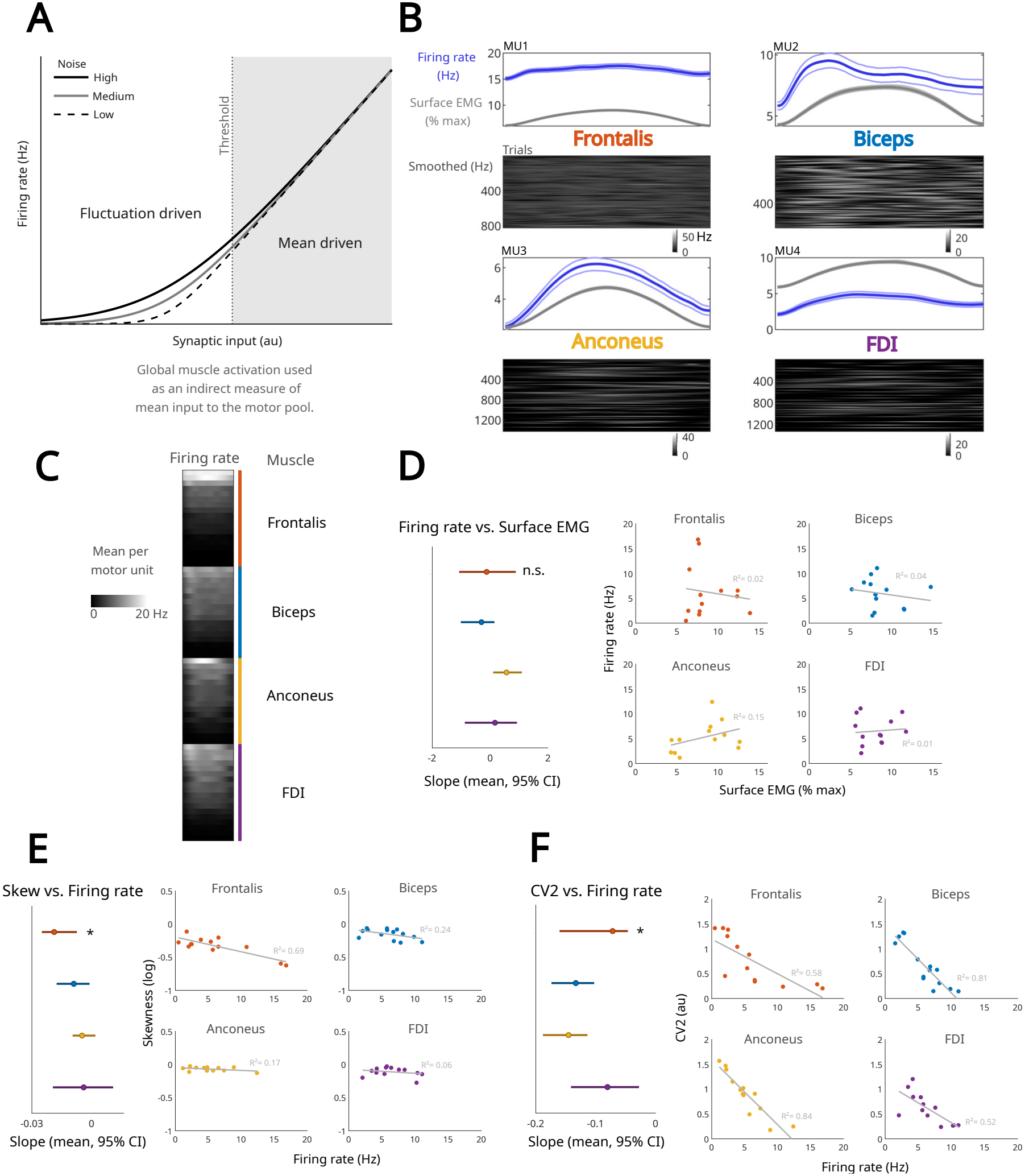
Firing regime differences across muscles based on voluntary activation curves. **A**, firing rate relationships were measured from changes of global muscle activation from the identified surface EMG curves, see figure 1c. Synaptic input below action potential threshold relates to a nonlinear input-output relationship (i.e., fluctuation driven, subprimary range) whereas higher synaptic input above threshold relates to a symmetric input-output relationship (i.e., mean driven, primary range). Adapted from Peterson et al.^23^ **B**, peri-event time histograms of motor unit activity locked to increases and decreases in whole muscle activation. Motor units here are the same as in Fig 1b,c (n = 1 participant). Firing rate and surface EMG curves are the mean with bootstrapped 95% confidence intervals. Trials show the mean firing rate across surface EMG curves. See supplemental figure 1 for data across trials per MU for this participant. **C**, mean firing rate from peri-event time histograms across all motor units, sorted by mean rate for each muscle. **D**, relationship between mean surface EMG and firing rates across motor units for each muscle (multiple linear regression, no difference in slopes across muscles, p > 0.23). **E**, mean firing rates and firing rate skewness across motor units for each muscle (multiple linear regression, * difference in slopes between the frontalis and anconeus, p = 0.02, and frontalis and FDI, p = 0.04). **F**, mean firing rates and CV2 (i.e., a local measure of firing rate irregularity) across motor units for each muscle (multiple linear regression, * difference in slopes between the frontalis and biceps, p = 0.02, and frontalis and anconeus, p = 0.02). D-E, in forest plots, the linear slope 95% confidence intervals are based on bootstrapping. In the scatterplots, each dot represents the ensemble average across trials for each motor unit. Grey lines show the linear regression fit; 13 motor units in the frontalis muscle, n = 9 participants; 14 motor units in the biceps muscle, n = 10 participants; 13 motor units in the anconeus muscle, n = 10 participants; 13 motor units in the first dorsal interosseous muscle, n = 10 participants. Firing rates included for calculations were from the changes of whole muscle activation from the surface EMG curves.

In the fluctuation-driven regime (analogous to the subprimary range), input is mainly below membrane threshold, and the input-output relationship is more sensitive to transient changes in input^23,25,31^. Furthermore, a noisy membrane threshold is shown to have minimal effect on this regime^23^. In contrast, the mean-driven regime (analogous to the primary range) operates above the membrane threshold and has a linearized input-output relationship^23,26,29,39^. By isolating motor unit spiking during the surface EMG detected activation curves, in which increases and decreases of whole muscle activation are relative to the steady-state baseline, we found diverse modulation responses across brainstem- and spinal-innervated muscles (Fig 2b,c).

Across motor units recorded, the relationship between surface EMG and firing rate was not different across muscles, corresponding to lower threshold motor unit activity that was <20 Hz (Fig 2d). Overall, compared with other muscles, the frontalis had the greatest firing variance from a similar range of voluntary activation among muscles. From prior modeling and spinal recordings^23,24^, increasing synaptic input results in a more negatively skewed firing rate distribution, representing a neuron operating in a mean-driven regime^23^. We confirm this relationship also occurs in brainstem and spinal motor units (Fig 2e), and found that the frontalis spent more time in the mean-driven regime than the spinal-innervated muscles.

A further robust metric to indicate the status of the firing regime is firing irregularity on a local time scale, termed CV2 (named from a modification of the coefficient of variation). A lower CV2 indicates less irregularity with synaptic input above membrane threshold for action potentials, as compared to a higher CV2 (generally above 0.5), which indicates a neuron operating in the fluctuation-driven regime with synaptic input below threshold^23^. We calculated CV2 across motor units for each muscle, and found that CV2 values spanned between 0.1 and 1.6 when locked to surface EMG curves (example curves, Fig 2b). In comparing the relationship of CV2 across muscles, we found that the frontalis operated with both regimes across motor units that had lower firing rates, as compared to spinal muscles (Fig 2f), indicating more firing regime flexibility in how frontalis motor neurons respond to synaptic input during voluntary activation. However, we found that the FDI muscle (spinal) had a similar slope relationship to the frontalis, possibly supporting prior work on flexible neural control of the hand as unique from other limb muscles^40–43^.

### Comparing firing regimes across brainstem and spinal motor pools

In comparing firing regimes between brainstem and spinal muscles (Fig 2), it was surprising that relationships were similar across upper limb muscles despite their different functional properties (Table 1). Therefore, to understand how control of the brainstem motor pool differs from a lower limb muscle, with leg muscles presumably having greater reliance on reflex and spinal control apart from motor cortex^44,45^, we used high-density surface EMG arrays placed over the right aspect of the frontalis and over the right tibialis anterior muscles during the same task (Fig 3a). Because contraction intensity was low, we recorded a high number of surface potentials with minimal signal complexity. This provided greater confidence in the accuracy of decomposed motor unit spike times (Fig 3b). The yield of motor units per session was lower in the frontalis muscle (range, 3-10) as compared to the tibialis anterior (range, 9-51), and this was likely due to anatomical differences in the number of active units that we could record from. Furthermore, despite recording from female participants (n = 3) we were unable to reliably decompose motor units in their frontalis and tibialis anterior muscles and therefore, the data were not included.

**Table 1.** Brief summary characteristics of recorded muscles.

| **Muscle** | **Innervation** | **Function** |
|---|---|---|
| Frontalis | Brainstem, cranial nerve 7, motor nuclei in the pons | Facial expressions that elevate the eyebrows, wrinkle the forehead, and pull the scalp forward |
| Biceps brachii | Spinal, musculocutaneous nerve, motor nuclei in C5-6 | Supination, elbow flexion and shoulder flexion |
| Anconeus | Spinal, radial nerve, C7 | Elbow extension and elbow stabilization |
| First dorsal interosseous (FDI) | Spinal, deep branch of the ulnar nerve, C8-T1 | Dexterous hand control that flexes and abducts the first digit |
| Tibialis anterior (TA) | Spinal, deep fibular nerve, L4-5, and S1 | Ankle dorsiflexion and inversion |

**Figure 3:**
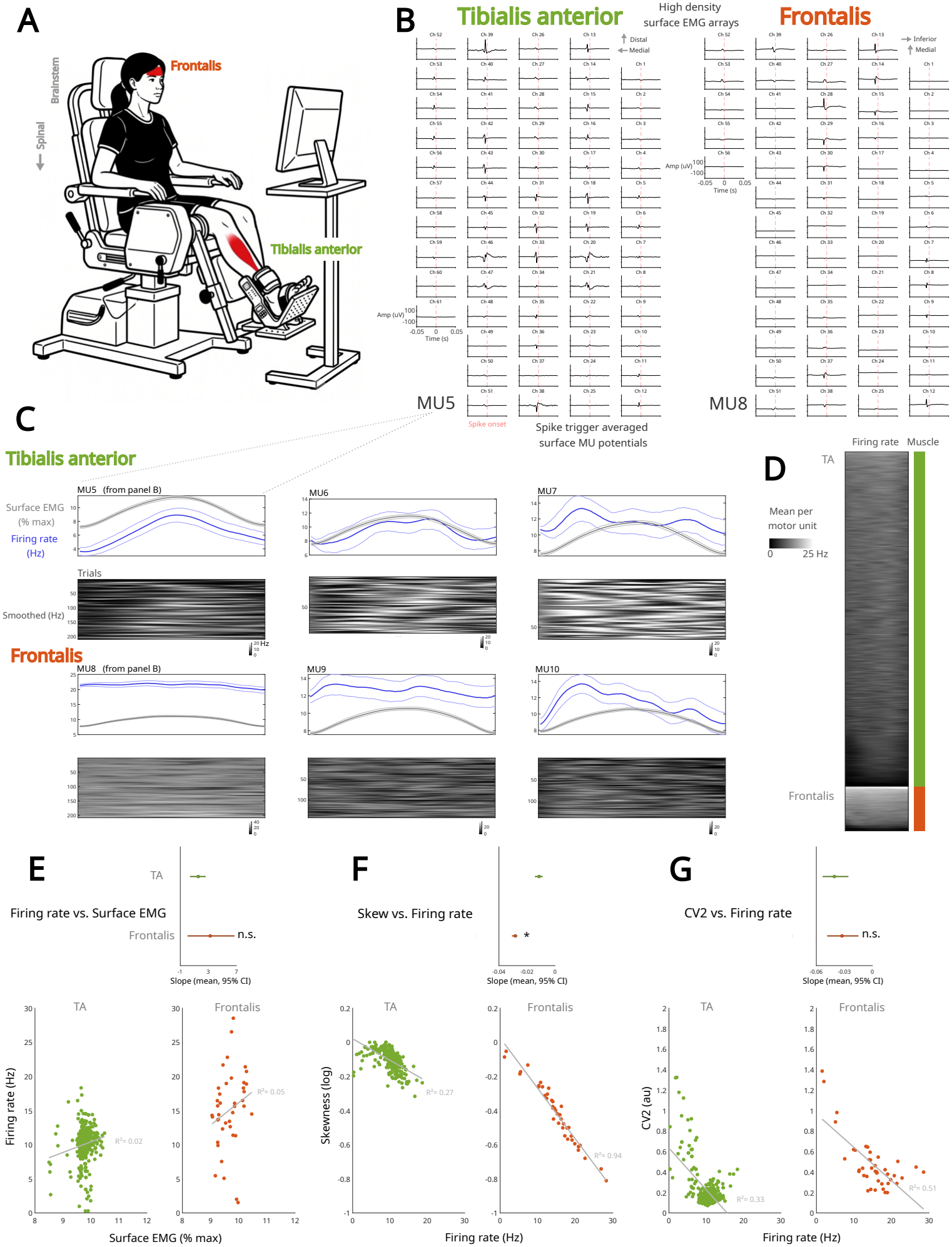
Voluntary activation of facial and lower limb muscles. **A**, cartoon showing the participant seated in a dynamometer with real-time visual feedback of muscle surface EMG. We recorded from the right aspect of the frontalis muscle (above the right eye), and from the TA on the right leg. Each muscle was recorded independently as the participant attempted to maintain a near-constant static contraction for ~2 min that was between 8 and 12% of maximum muscle activation. **B**, motor unit spike trigger average waveforms for each muscle across high-density surface EMG arrays. MU5 was recorded in the TA across 61 channels with 1272 spikes identified (n = 1 participant), and MU8 was recorded in the frontalis across 56 channels with 3674 spikes identified (n = 1 participant). Mean voltage traces are 100 ms, with 95% confidence intervals based on bootstrapping. **C**, peri-event time histograms of motor unit activity locked to increases and decreases in whole-muscle activation (n = 2 participants). MU5 and MU8 are linked to panel B. Firing rate and surface EMG curves are the mean with bootstrapped 95% confidence intervals. Trials show the mean firing rate across surface EMG curves. **D**, mean firing rate from peri-event time histograms across all motor units, sorted by mean rate for each muscle. **E**, relationship between mean surface EMG and firing rates across motor units for each muscle (multiple linear regression, no difference in slopes, p = 0.23). **F**, mean firing rates and firing rate skewness across motor units for each muscle (multiple linear regression, * difference in slopes, p < 0.001). **G**, mean firing rates and CV2 across motor units for each muscle (multiple linear regression, no difference in slopes, p = 0.14). D-F, in forest plots, the linear slope 95% confidence intervals are based on bootstrapping. In the scatterplots, each dot represents the ensemble average across trials for each motor unit. Grey lines show the linear regression fit; 341 motor units in the TA muscle, n = 4 participants; 41 motor units in the frontalis muscle, n = 4 participants. Firing rates included for calculations were from the identified changes of whole muscle activation from the surface EMG curves.

Similar to the intramuscular recordings (Figs 1-2), again we identified surface EMG curves representing increases and decreases of volitional input to the motor pool, and visualized the relationship of firing rates across motor units (Fig 3c,d). Motor units identified in the frontalis from surface decomposition had responses similar to those from intramuscular recordings in different participants. From the large sample of motor units, the frontalis muscle had higher firing rates as compared to the tibialis anterior within a similar range of relative voluntary effort (Fig 3e), indicating a greater role of rate coding than recruitment in the brainstem muscle.

In assessing measures to estimate the firing regime used by each muscle, we found that skewness had a strong linear relationship to increasing firing rate in the frontalis as compared to the spinally innervated leg muscle (Fig 3f), indicating more time spent in the mean-driven regime. Across motor units, this shows that outputs in the frontalis are more representative of the input, corresponding to a form of voluntary control that may not need further processing by intrinsic neuronal mechanisms (such as persistent inward currents or premotor interneurons)^2,11,35^. In using another metric to assess the firing regime, we found that the average CV2 was different between muscles but the linear relationship to firing rates was not (Fig 3g). In the spinal-innervated muscle, at ~9 Hz, a transition occurs when the regime becomes mean-driven, whereas in the frontalis, CV2 has more variability across lower and higher firing rate motor units; and in agreement with the intramuscular recordings, we show that the brainstem-innervated muscle can operate across both the fluctuation- and mean-driven regimes.

### More flexible neural control in the brainstem-compared to spinal-innervated muscle

We found that frontalis motor units displayed properties of more flexible voluntary control as compared to tibialis anterior motor units. Based on our anatomical hypothesis for flexible control, a muscle with more direct cortical input should have greater neural flexibility. A single aspect of the frontalis muscle (for example, above one eye) receives corticobulbar tract inputs from both hemispheres^1,2^, whereas the tibialis anterior is predominantly supplied from indirect inputs from the corticospinal tract in the contralateral hemisphere^46,47^, along with many other descending tracts and reflex pathways (Fig 4a).

**Figure 4:**
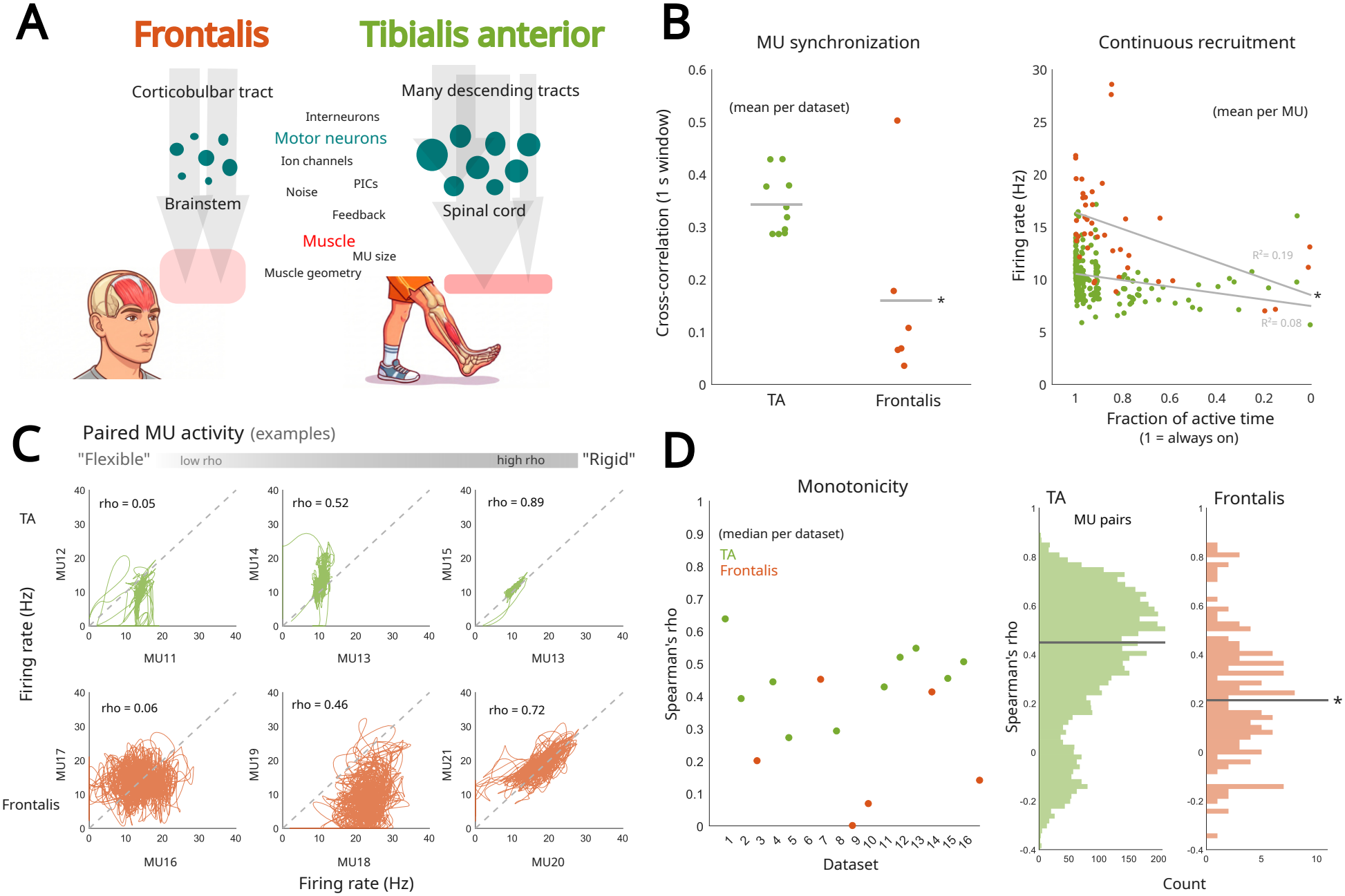
Control of the motor pool differs between brainstem-versus spinally-innervated muscle. **A**, cartoon and schematic of some important differences between muscles. **B**, leftmost, pairwise firing rate cross-correlation across motor units for each muscle with each dot representing the mean per session; the grey horizontal line shows the ensemble mean, * difference in lines, Wilcoxon rank-sum test, p = 0.0312. **B**, rightmost, relationship between the fraction of activity during the session and firing rate across motor units. Grey lines show the linear fit for each muscle (multiple linear regression, * difference in slopes, p = 0.001). **C**, firing rate relationship of monotonicity between example motor units that were simultaneously recorded. The Spearman’s correlation coefficient (rho) is displayed for each pair. **D**, leftmost, pairwise relationship of monotonicity across motor units for each muscle, with each dot representing the median Spearman’s coefficient per session. **D**, rightmost, pairwise Spearman’s correlations across all motor units recorded simultaneously within each session. The horizontal grey line shows the ensemble average per muscle across datasets (TA rho = 0.45, frontalis rho = 0.21; * difference in lines, Wilcoxon rank-sum test, p = 0.016). B-D, 341 motor units in the TA muscle, n = 4 participants; 40 motor units in the frontalis muscle (one less than Figure 3), n = 4 participants.

Because recordings were sustained for long durations, we were able to capture more complex behavior outside of only aligning firing rates to the surface EMG curves (see Figures 2 and 3). This provided the ability to include longer periods of sustained voluntary activity, ramp contractions and brief rest. From pairwise comparisons of motor unit firing within each session, the frontalis had lower synchronized activity than the tibialis anterior (Fig 4b, left). To assess whether this was due to a lack of continuous activity explained by differences in motor unit recruitment, we compared the relationship of firing rate to the fraction of active time (Fig 4b, right). We found that the majority of frontalis motor units were indeed active throughout the session (Fig 4b, right), and inactivity was unlikely to explain differences in synchronization between muscles.

Subsequently, we used an approach to quantitatively assess whether motor unit activity can occur with features of independent voluntary drive^15,21^, which would indicate a greater role for flexible control. This involved calculating the strength of monotonicity across motor unit pairs within each session, and examples are shown for each muscle (Fig 4c). Under the assumption of distributed neural drive to the motor pool, rigid control would satisfy positive monotonicity between motor unit pairs within a similar voluntary activation range, whereas violations would support independent drive and therefore flexible control. Overall, we found that the frontalis muscle had greater independent control across motor units than the tibialis anterior^15,16^, providing evidence that brainstem voluntary control is more flexible as compared to the spinal muscle (Fig 4d). However, when plotting all pairwise relationships, we also found that the tibialis anterior had a small fraction of motor units that displayed features of flexible control (Fig 4d, right). In these cases, the tibialis anterior pairs were largely influenced by recruitment threshold differences, with one motor unit being recruited in a mean-driven regime as compared to another motor unit that resided largely below recruitment threshold, existing in the fluctuation-driven regime, suggesting motor unit functional clusters within the same muscle^15,48^.

## Discussion

Everyday movements happen along a natural continuum of low-effort activity. During these voluntary contractions, we found that spinal-innervated muscles displayed more features of rigid control when examining measures of the firing regime and paired motor unit activity. In contrast, the brainstem-innervated muscle had properties of more flexible control. These findings build on prior work^21,40–43^, and our results support a theory of neuromotor flexibility based on anatomical constraints that we hypothesize is dependent on the degree to which muscle can be accessed by independent elements of the nervous system.

In humans, some facial muscles, like the frontalis, are supplied by brainstem motor nuclei that are under near-equal control from both cortical hemispheres^1–6,49–51^, whereas spinal motor pools receive the majority of their cortical commands from the contralateral hemisphere^52–54^ and many subcortical pathways^55,56^. Because the pons, which houses brainstem motor nuclei, is a relatively newer evolutionary development as compared to the spinal cord^57,58^, it remains likely that more recent mammalian circuits that involve greater inputs from motor cortices achieved more flexible-like control of muscle^7,59,60^. Likewise, greater excitability of brainstem as compared to spinal motor neurons^10^ would presumably reduce the burden of excitatory synaptic inputs to achieve relative levels of activation. In the framing of rigid versus flexible neural control, there are many additional descending spinal tracts and reflex pathways that play important roles^61,62^. For example, facial muscles are under different reflex circuits than spinal-innervated muscles, with the frontalis having minimal corpuscular afferent neuronal structures^63^, whereas limb muscles receive strong inputs from antagonist muscle reflexes^64–66^. By contrast, some muscles, like the superior trapezius, receive both brainstem and spinal innervation^67^, but other muscles like the deltoid, which appear to be controlled in a flexible way^21^, receive spinal innervation across seven different compartments^68^. In determining the capacity of a muscle for flexible control based on cortical involvement, the complexity of the whole-muscle force profile becomes less important because there are numerous ways motor pools can be activated to produce a similar movement pattern^21,69,70^. Instead, greater emphasis on how neural elements work together to achieve different aspects of movement would broaden understanding about what features of control are accessed across what level of the sensorimotor hierarchy.

The strength of distributed synaptic inputs across the motor neuron pool influences how motor units are recruited and rate coded. Although recruitment corresponds to the necessary activation of muscle, it is rate coding that enables force profiles to be shaped and continuously modified. Rate coding has been described in relation to three input-dependent firing regimes, which are defined as sub-primary (i.e., fluctuation-driven)^23,25,26,31,32,36^, primary (i.e., mean-driven)^23,28,29,71^ and the secondary range^28,33^. We show in human brainstem and spinal muscles that both fluctuation- and mean-driven firing regimes are used to maintain low-effort voluntary contractions, in agreement with population recordings of evoked reflexes in the semi-intact turtle spinal cord^23^. These two firing regimes correspond to a diversity of flexible and rigid control modes within and between differently innervated motor pools. Because brainstem and spinal networks use both regimes across their motor units (i.e., motor outputs or readouts), this dual-regime utilization is shown to provide necessary properties of network stability while being sensitive to new synaptic input^24,72–74^. However, we show that the brainstem had a greater proportion of higher firing rate motor units in the fluctuation-driven regime, supporting a wider range of sensitivity to voluntary input^11^.

The utility of a muscle using both firing regimes can be understood in how contractile forces are generated and maintained. Conventionally, lower threshold motor units are shown to have greater utility of a fluctuation-driven regime because the afterhyperpolarization and contractile relaxation properties are longer^25,31,32^ as compared to higher threshold motor units^75,76^. But, whole-muscle mechanics during movement are also an important consideration, because facial muscles would use more viscoelastic forces as compared to limb muscles that would require complex series-elastic and inertial forces^77–80^. For example, the mean-driven regime would be useful to generate or sustain larger forces whereas the fluctuation-regime could maintain a weaker tone or make small force adjustments. Another exception of distributed inputs to the motor pool would be cortico-motoneuronal cells, which provide the anatomical substrate for flexible neural control. However, it remains unclear whether they are more prominent in human brainstem versus spinal muscles (e.g., intrinsic hand), or whether there is bias for smaller versus larger motor units^40–43^. Regarding dexterous control, evidence in humans shows that intrinsic muscles of the hand are innervated with far fewer motor axons than previously expected, having ~1700 motor axons across ~20 muscles^8^. Expectedly, the ~42 muscles found in the face may be similarly controlled by relatively few motor axons, and under the framework of flexible control, independent access by cortex would expand the functional repertoire of effectors^59^ when limited by the number of motor units.

A limitation is that we were unable to directly record motor neuron membrane thresholds, but instead we inferred the firing regimes based on measures that have been validated using computational and spinal recording preparations^23,24^. Furthermore, we did not characterize motor unit size (which would have provided a secondary measure to understand if recruitment violations were size-based); instead, we relied on firing rate and recruitment order comparisons across motor unit pairs^14,15,21^. Finally, our voluntary task was simple, and for practical reasons, we chose low-effort and sustained contractions to increase our confidence in isolating motor unit potentials across different muscles. Therefore, the voluntary dynamic range in this task was relatively small and influences from biological noise and measurement errors in missing action potentials should be taken into account. Yet, more complex movements are desired because they are likely to yield more striking neural control differences based on muscle innervation properties.

In comparing brainstem and spinal pools, we tested different features of motor unit control from several muscles across the neuromotor axis. We show that the brainstem-innervated muscle has more flexible neural control properties at both the resolution of single motor units and across the motor pool in comparison to several spinally innervated muscles we tested. Grounding the hypothesis for flexible neural control within anatomical constraints, it is conceivable that evolutionarily newer circuits, like those that innervate the frontalis^7,81,82^, are accessed by more independent cortical elements that can generate nuanced movements, allowing animals to raise their eyebrows quizzically or with surprise!

## Methods

### Participants

For intramuscular recordings, ten healthy adults (8 male and 2 female) provided informed consent. All procedures were approved by the local research ethics board for human experimentation, and the study occurred at the University of Western Ontario, London, Ontario, Canada, in the laboratory of C.L. Rice. Each participant was involved in 2-4 experimental sessions that each lasted between 1 and 3 hours. The frontalis muscle data from one male participant were excluded from analysis because no motor units could be identified.

For high density surface recordings, seven healthy adults (4 male and 3 female) provided informed consent. All procedures were approved by the local research ethics board for human experimentation, and the study occurred at Northwestern University, Chicago, Illinois, USA, in the laboratory of C.J. Heckman. Each participant was involved in 1-2 experimental sessions that lasted between 1 and 3 hours. Three female participants were excluded from the results because we were not able to decompose motor units from their frontalis and tibialis anterior muscles.

### Voluntary task with intramuscular recordings

In the seated position, participants made voluntary contractions focusing on one muscle at a time. Each muscle was activated between 1 and 18% of the maximum voluntary contraction (MVC) for a duration of 3-12 minutes, with real-time feedback of surface EMG. Participants slowly contracted the desired muscle, and were instructed to slowly increase and decrease muscle activation based around the firing threshold of single motor units, or a set activation percentage. Joint positions and facial expressions were kept relatively static throughout the recordings, with brief periods of rest (1-5 seconds) throughout.

### Intramuscular EMG

Two approaches were required for single MU recordings, dependent on the muscle being studied. For the frontalis muscle, we needed to manipulate a monopolar needle electrode (insulated tungsten wire, 250 µm diameter, 1 µm tip with a 30 degree taper angle, Fredrick Haer) that was inserted into a transverse wrinkle of the frontalis muscle, approximately 0.5-2 cm above the eyebrow. The needle was sterilized using an autoclave, and was inserted at an acute angle of 10-30 degrees and was advanced gently through superficial aspects of the muscle, and parallel to the selected transverse wrinkle (i.e., cutaneous rhytid or forehead line) until a stable motor unit was identified. Real time visual and audio feedback provided to the investigator confirmed single MU recordings. A reference electrode (2.5 × 2.5 cm, silver Mactrode electrodes, GE Medical Systems, Milwaukee, WI) was placed over the zygomatic arch on the side of the face, and this recording pair was grounded on the clavicle^11^. For upper limb muscles, bi-polar electrode recordings were used. For each bi-polar electrode, the fine wire (0.102 mm diameter, insulated stainless steel wire, California Fine Wire Company, Grover Beach, CA) was cut perpendicular and bent with fl0.5 cm hooks prior to being threaded into the hypodermic needle (27 gauge), autoclaved, and then inserted into the belly of each muscle. Across muscles, the common ground electrode was placed on the clavicle. For both intramuscular recording methods, each recording channel was amplified 100-200x, hardware filtered between 10 Hz and 10 kHz (Neurolog, Welwyn City, UK) and sampled at 25 kHz (Power1401, Cambridge Electronic Design, Cambridge, UK). During off-line analysis in Spike2 software (version 7, Cambridge Electronics Design, Cambridge), intramuscular EMG was digitally high pass filtered at 0.5 kHz using a 2^nd^ order infinite impulse response Butterworth filter. Motor unit spike times were then identified based on voltage thresholding and waveform template matching. For each muscle, the intramuscular and surface EMG recordings were simultaneously recorded and synchronized.

### Monopolar surface EMG

Surface EMG was recorded in a monopolar configuration in the frontalis muscle and upper limb during intramuscular recordings. For the frontalis muscle, the active electrode was placed ~5 cm above the right eye, and the reference was placed on the frontal bone just lateral to the muscle (silver Mactrode electrodes, GE Medical Systems, Milwaukee, WI). For each upper limb muscle, the active electrode (0.9 cm diameter, silver chloride H59P, Kendall, Covidien, Massachusetts) was placed on the midpoint location at the muscle belly and the reference was placed on the distal tendon. Finally, a common ground electrode was placed on the right clavicle. Each channel was hardware filtered between 20 Hz and 2 kHz, amplified 200x (Neurolog) and was sampled at 5 kHz (Power1401). Real time visual feedback of the surface EMG signal for each muscle was provided during each experiment, and was full wave rectified and smoothed with a 0.1 s time constant. To calibrate the voluntary contraction intensity during experiments, the maximum amplitude during a 5 second maximum voluntary effort was used to represent 100% activation. During high-density surface recordings, surface EMG for feedback was from one of the channels with signal-to-noise above the 95th percentile.

### Voluntary task with high-density surface recordings

Participants were randomized to perform either dorsiflexion or frontalis contractions first. Prior to testing each muscle, participants completed a familiarization block where they were instructed on proper contraction technique and performed 5-10 practice trials until proficient. Next, participants performed three MVCs with 60 seconds rest between each attempt. If the rectified EMG amplitude of the final MVC trial was greater than that of the preceding attempts, additional MVC trials were performed as needed. Verbal encouragement was given to participants during all MVC trials to ensure maximal effort, and for the frontalis muscle participants were cued to “raise their eyebrows” with maximal effort. The peak rectified EMG amplitude achieved during the MVC was used for the normalization of subsequent trials. For submaximal trials, participants completed 2 minute contraction holds at 10% MVC for each muscle. Trials were repeated as necessary until two successful relatively steady holds were achieved. Participants received real-time visual feedback of the EMG from the target muscle (tibialis anterior or frontalis) during each trial.

### High-density surface EMG

Prior to placing electrodes, excess hair was shaved from the area and the skin was lightly abraded with skin prep gel (Spes Medica, Everi, Genoa, Italy). For the leg, a single high-density surface EMG (HDsEMG) array (64 electrodes, 13 × 5, 8 mm inter-electrode distance, GR08MM1305, OT Bioelettronica, Inc., Turin, Italy) was placed vertically on the tibialis anterior muscle of the right leg. A reference electrode strap was secured around the medial and lateral malleolus on the right ankle. In the dynamometer, the ankle was positioned at 85 degrees, so that the dorsiflexor muscles were slightly shortened as compared to the 90-degree neutral position. For the forehead, a single HDsEMG array (64 electrodes, 13 × 5, 4 mm inter-electrode distance, GR04MM1305, OT Bioelettronica, Inc., Turin, Italy) was placed horizontally on the right frontalis muscle (above the right eye). A reference electrode was placed on the clavicle. HDsEMG data were sampled at 2048 Hz per channel, amplified x150, band-pass filtered 10-500 Hz, and collected in differential mode using a Quattrocentro signal amplifier (OT Bioelettronica, Inc.).

Prior to collecting experimental data, real-time HD-sEMG recordings were checked visually to ensure high signal-to-noise ratios. During experiments, real-time visual feedback of a single channel was amplified (x16) using the OTbio+ software from the analog output feature^83^. The EMG channel was chosen based on its central location on the electrode array and its signal-to-noise ratio. This EMG channel was routed to a National Instruments data acquisition board (NI-USB-6218) and processed in real time using a custom MATLAB script to generate visual feedback on a TV monitor. Prior to displaying the feedback, the EMG was lowpass filtered (1.8 Hz), corrected for resting baseline activity, and averaged over 60 ms intervals.

#### Motor unit decomposition from high-density surface EMG

Off-line, each HDsEMG channel was manually inspected, and channels with artifacts or non-physiological noise were removed. Motor unit spike trains were decomposed from the remaining HDsEMG channels using a convolutive blind-source separation method^84^ with a silhouette threshold of 0.85. Using a custom graphical user interface in MATLAB, each MU spike train was visually inspected, and edited manually by a trained technician to correct minor errors made by the decomposition algorithm, using similar techniques described in recent studies^83,85,86^.

#### Analysis

##### Identifying peak curves from surface EMG

To estimate changes of whole muscle activation, surface EMG was used as a proxy for the mean synaptic drive of volitional input to each motor pool. Surface EMG was bandpass filtered 20-500 Hz, rectified and smoothed with a 100 ms Gaussian. Surface EMG was then normalized to a percentage from the 5 second plateau of the maximum effort contraction (i.e., MVC) at the same joint position. Temporary peaks from the normalized smoothed surface EMG were then identified to represent the times of mean drive increases and decreases throughout each muscle recording. Each surface EMG curve was identified by finding local peaks throughout the recording that were required to occur at least 0.5 seconds apart, have a minimum relative amplitude change of 0.5%, a minimum peak width duration of 100 ms and a minimum peak prominence of 1.2% from adjacent peaks. We also performed analyses requiring a minimum peak width duration of 250 ms, and found the same relationships within and across muscles (data not shown) as presented in the results.

##### Firing rates

Across identified motor unit spike times for each session, the firing rate time series was computed using Gaussian smoothing with a 100 ms kernel. Because voluntary contraction intensity was low, instantaneous firing rates >150 Hz were excluded prior to smoothing. Based on the timing of identified peak curves from the surface EMG, smoothed firing rates and surface EMG were aligned and time warped by linear interpolation for generating peri event time histograms (PETHs).

##### Skewness of distribution

We used skewness (third moment) as a measure of asymmetry of the firing rate distribution around its mean. In some muscles individual surface EMG curves contained too few spikes for reliable skewness estimation (less than 4), therefore, for each motor unit, surface EMG curves (i.e., trials) were grouped into 15 equal-count quantile bins according to the mean instantaneous firing rate. We computed the bias-corrected sample skewness (equivalent to the G1 adjusted Fisher-Pearson estimator) from the firing rate distribution that was adjusted on a logarithmic scale^23,24^.

##### Irregularity of spiking activity

For each motor unit, we calculated a time dependent metric of firing rate irregularity (CV2), calculated across the time series of spiking activity for each session^23^. CV2 was defined by pairs of adjacent inter-spike intervals, where CV2 =1, for a Poisson process (i.e., fluctuation-driven firing regime), and CV2 = 0, for a regular process (i.e., mean-driven firing regime).

##### Cross-correlation and continuous recruitment metrics

From motor unit spike times decomposed using high density surface EMG arrays, firing rates were smoothed with a 0.5 s Hanning window (Fig 4b). For motor units that were active across the session, we calculated pairwise cross-correlations with a sliding 1 second window. This resulted in a distribution of cross-correlation coefficients for each session. For a measure of continuous recruitment, the fraction of active time (i.e., time spent recruited) was calculated based on how long the motor unit was above a firing rate threshold of 2 Hz.

##### Estimate of monotonicity between motor units

Spearman rank correlations were computed across the continuous firing rate time series for all motor unit pairs within each session^12,14,15,21^. Firing rates were smoothed with a 0.5 s Hanning window. Throughout the session, we restricted the analysis to time bins where at least one motor unit in the pair maintained a firing rate greater than 2 Hz.

##### Linear regression

Linear regression was used with an interaction term for muscle group (Figs 2d-e, 3e-g, and 4b). For comparisons across muscles, an analysis of variance was performed on the fitted model. To determine significance between muscle slopes, we iterated through all pairwise comparisons refitting the linear model. To assess the goodness-of-fit, the coefficient of determination from ordinary least squares is reported within respective figures.

## Data and code availability

Data and code can be accessed by reasonable request from the corresponding author. Processed data used in the results will be made available in a public repository upon publication.

Inkscape version 1.4.4 and Google Gemini 4, a large language model, were used to edit cartoons in figures 1a, 3a, and 4a. Also, Gemini was used for editing code, with all code modifications tested by human review.

## Contributions

E. Kirk was involved with all aspects of the study. S. Jenz, J. Beauchamp, A. Benedetto, G. Pearcey, and C.J. Heckman were involved with HD-sEMG experiments. C. Rice was involved with intramuscular experiments. All authors were involved with conceptualization and editing the manuscript. The authors declare that they have no competing financial interests or personal relationships that could have appeared to influence the work reported in this paper.

## Acknowledgements

We thank Dr. T. Doherty for their insight and training on frontalis muscle recordings. Likewise, we acknowledge Drs. A. Zero, J. Fanous and B. Sauerbrei for helpful discussions. Finally, we thank former lab members in London and Chicago, and all participants for their involvement.

---

**Supplement Figure 1:**
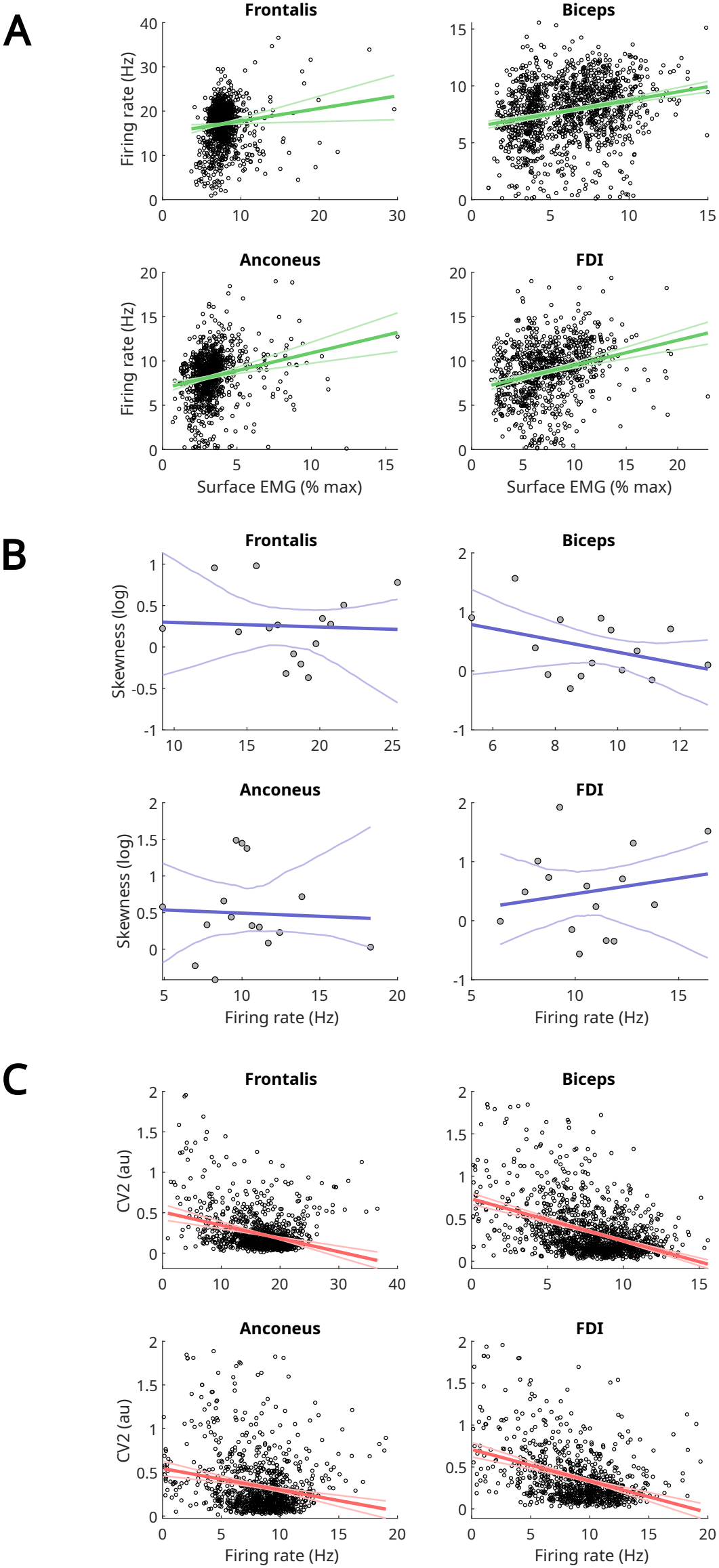
Average values across voluntary activation curves for motor units highlighted in figures 1 and 2b (n = 1 participant). **A**, relationship between mean surface EMG and firing rates for each muscle. **B**, mean firing rates and firing rate skewness for each muscle. For each motor unit, surface EMG curves (i.e., trials) were grouped into 15 equal-count quantile bins according to the mean instantaneous firing rate. **C**, mean firing rates and CV2 for each muscle. For A and C, each dot represents the average value per trial (i.e., surface EMG curve) for each motor unit. Lines show the linear regression fit and the 95% confidence intervals are based on bootstrapping.

